# Ethanol alters mechanosensory habituation in *C. elegans* by way of the BK potassium channel through a novel mechanism

**DOI:** 10.1101/2024.11.22.624824

**Authors:** Nikolas Kokan, Conny Lin, Alvaro Luna, Joani Viliunas, Catharine H. Rankin

## Abstract

In this research, we investigated how alcohol modulates the simplest form of learning, habituation, in *Caenorhabditis elegans*. We used our high throughput Multi-Worm Tracker to conduct a large scale study of more than 21,000 wild-type worms to assess the effects of different doses of alcohol on habituation of the well-characterized tap withdrawal response. We found that the effect of alcohol on habituation of the reversal response to repeated mechanosensory stimuli (taps) differed depending on the component of the reversal response assessed. Furthermore, we discovered that alcohol shifted the dominant response to tap from a backward reversal to a brief forward movement. Because the large conductance potassium (BK) channel has been shown to be important for alcohol’s effects on behaviour in a variety of organisms, including *C. elegans*, we investigated whether the *C. elegans* BK channel ortholog, SLO-1, mediated the effects of alcohol on habituation. We tested several different strains of worms with mutations in *slo-1* along with wild-type controls; null mutations in *slo-1* made animals resistant to alcohol induced changes in learning. However, a mutation in the putative ethanol binding site on SLO-1 did not disrupt ethanol’s impact on habituation. Finally, by degrading SLO-1 in different parts of the nervous system we found that SLO-1’s function in ethanol’s impact on habituation is likely distributed throughout the neural circuit that responds to tap. Based on these results, our main conclusions are 1) ethanol is not a general facilitator or inhibitor of habituation but rather a complex modulator, 2) SLO-1 is required for ethanol’s effect on habituation, 3) ethanol is interacting (directly or indirectly) with SLO-1 through a novel unidentified mechanism to influence response plasticity.

## Introduction

Although alcohol alters many behavioral and cognitive functions including physical coordination, attention, and learning and memory (1,2) alcohol’s effects on the simplest form of learning, habituation, are not well understood. Habituation is defined as a decremented response to repeated stimuli that is not due to fatigue or sensory adaptation (3). Habituation is highly conserved across the animal kingdom and is thought to be a foundation of selective attention, which allows animals to filter out unimportant stimuli and focus on stimuli that have biological consequences. Several neuropsychiatric disorders have been associated with altered habituation (for a review see McDiarmid *et al*., (4)). For example, patients diagnosed with schizophrenia have slower habituation to sensory stimuli, which is hypothesized to be an underlying reason for their over-reactivity to stimuli (5–11). Although habituation’s role in alcohol and substance use disorders has not received the same attention as other psychiatric disorders, there is evidence that abnormal habituation may underly cognitive dysfunctions associated with drug addiction (12).

The existing literature on the relationship between alcohol intoxication and habituation is limited and contradictory. The first report on acute alcohol intoxication and habituation in humans investigated habituation of post-rotatory nystagmus (rapid involuntary movement of the eyes) in fighter pilots, ballet dancers, and figure skaters (13). This study found that these subjects, known to have exceptional post-rotatory nystagmus habituation, showed no habituation when intoxicated. However, subsequent studies in humans and various other animals showed inconsistent results. Some studies found alcohol inhibited habituation (frog: 13, cichlid fish: 14, isolated frog spinal cord: 15, mice: 16), similar to the impaired habituation of post-rotary nystagmus (13,18), while other studies found alcohol facilitated habituation (cichlid fish: 14, isolated frog spinal cord: 18, human: 19). Although most of the previous studies measured only a single component of behaviour, a study by Peeke *et al*. (15), found that alcohol had opposite effects on habituation of different aspects of aggressive displays in male cichlid fish. Alcohol-exposed fish showed faster habituation of response frequency, but slower habituation of response duration. These data suggest that in the same animal alcohol can have divergent effects on different components of the same response and raised the possibility that alcohol may not globally inhibit or facilitate habituation.

In this paper, we used a well-characterized genetic animal model, the 1mm long nematode *Caenorhabditis elegans*, and high-throughput behavioral analyses to investigate alcohol’s effect on habituation. A mechanical tap to the side of their petri dish holding the worms will induce *C. elegans* to produce a reversal, which is a brief backward movement that has been called the “tap withdrawal response” (21). This tap withdrawal response has been well characterized (20–24, see review: 25). When a series of taps are given at a fixed inter-stimulus interval (ISI), worms respond less and less frequently to the tap stimuli and their responses get progressively smaller. In previous studies using our high throughput machine vision Multi-Worm Tracker apparatus, we discovered that different features of the response to tap (i.e., probability, duration, and speed) can be mediated by different genes (26,27). These data led to the conclusion that there is not a single mechanism of habituation, but that habituation of different response components can be mediated by different genes. The insights from this detailed analysis of tap habituation in *C. elegans* offer an excellent opportunity to clarify the effects of ethanol on habituation.

A variety of alcohol-related behaviors have also been studied in *C. elegans* (28–43). Ethanol has been shown to increase or decrease locomotor activity depending on the dose of ethanol exposure (28,44,45). Ethanol also inhibits egg-laying and increases muscle contractions when worms are on food (29,46). More complex alcohol-related behaviors such as state-dependent learning have also been reported in this organism (47).

A key contributor of alcohol’s effects on behavior is the BK channel, or the <u>B</u>ig conductance voltage- and calcium-gated Potassium(<u>K</u>+) channel. Outside of alcohol metabolic enzymes, the BK channel was the first protein whose physiological response to ethanol was linked to the behavioral effects of ethanol in the same animal. Notably, this study was carried out using *C. elegans* and since this first association of alcohol’s effect on behavior and the BK channel (29), the effect of alcohol on BK channels has been well-documented in a variety of organisms ranging from fruit flies to mammals (48–51). In *C. elegans*, genetic mapping of strains with strong resistance to ethanol showed that they all carried mutations in the BK channel ortholog *slo-1* (29). This resistance to ethanol was eliminated when wild-type copies of *slo-1* were restored in neurons but not in muscles (29). Furthermore, this study found that ethanol increased the SLO-1 current by increasing the frequency of channel opening (29). Gain of function *slo-1* mutants showed a crawling speed phenotype comparable to wild-type worms on ethanol, providing further support that hyperactivity of SLO-1 is responsible for ethanol’s inhibitory effect on crawling speed. In addition, the SLO-1 T381I substitution mutation, which is located 10-15 residues away from the SLO-1 RCK1 calcium sensor, resulted in strong resistance to ethanol’s effect on crawling speed and egg laying (52). The T381 amino acid residue is conserved in humans, mammals and flies and is part of an ethanol binding site on the BK channel that was later identified based on amino acid substitution, electrophysiology, and crystallography data (50,52–54). This ethanol binding site has been found to be important for *slo-1* mediated effects of ethanol on behavior in *C. elegans* (52).

The hypothesis of the study reported here is that ethanol is not a general facilitator or inhibitor of habituation; instead, it can have different effects on habituation of different components of a behavioral response. After first validating our approach by replicating the effect of ethanol on body curvature, we habituated animals to taps on and off ethanol. We found that ethanol had different effects on three different components of habituation to tap: response probability, duration and speed. Furthermore, using detailed behavioral analyses we found ethanol not only affected habituation of reversal responses to repeated taps, but also switched the dominant response from a reversal to a rapid forward movement. Because *slo-1* is known to be important for other behavioral effects of ethanol, we tested whether *slo-1* is also required for ethanol’s effects on habituation to repeated taps and found that the effect of ethanol on habituation was lost in *slo-1* null mutant worms. Surprisingly, the T381I region of SLO-1 that is part of an ethanol binding site on the BK channel was not important for ethanol’s effect on habituation, suggesting another site on SLO-1 is mediating its effect. By degrading SLO-1 specifically in the nervous system or the mechanosensory neurons we showed that ethanol’s effect on habituation occurred in the nervous system and not in muscles. These findings suggest that the function of SLO-1 in mediating ethanol sensitivity of response probability was most likely distributed throughout the neural circuit underlying the response to tap.

## Materials and Methods

### Worm culture

*C. elegans* stocks were grown and maintained on Nematode Growth Medium (NGM) plates with the *Escherichia coli* OP50 strain as a food source (55). N2 Bristol strain was used as the reference wild-type strain. N2, *slo-1(eg142)* BZ142 (backcrossed 2X), and *slo-1(js379)* NM1968 (backcrossed 5X) strains were obtained from the *C. elegans* Genetic Center (CGC). The *slo-1(gk602291)* JPS429 (backcrossed 6X) strain was a gift from Jon Pierce-Shimomura (52). The *slo-1(cim105[slo-1::GFP])* HKK796 *slo-1::GFP* strain and *slo-1*(*cim105*[*slo-1*::GFP]); cimSi1[*rgef-1p*::vhhGFP::*zif-1*::operon-linker::mCherry::*his-11*::*tbb-2* 3’UTR+cbr-*unc-119*(+)] HKK1165 neuron specific degraded strain were gifts from Dr. Hongkyun Kim’s laboratory (56). The strains with touch-neuron specific degradation of *slo-1::GFP* were made from crossing HKK796 and OD2984 [ltSi953[*mec-18p*::vhhGFP4::*zif-1*::operon-linker::mKate::*his-11*::*tbb-2* 3’UTR + Cbr-*unc-119*(+)] II; *unc-119*(ed3) III] which were obtained from the CGC.

### Culture age synchronization for behavioral testing

To obtain age-synchronized colonies of animals for testing, 12.5µl of bleach reagent composed of 1:1 Bleach, 5% solution of sodium hypochlorite and 1M NaOH, as described in (57), was pipetted onto a OP50 free area of a 5 cm NGM plate seeded with OP50. A number (at least 15) of gravid adult worms were placed in the bleach dot. Adults were bleach lysed and the eggs in their gonads were released; within a few hours the larval worms hatched from the eggs and crawled to the *E. coli* (57). The resulting colony was made up of 20-80 worms that were age matched within a range of 350 minutes, or 5-6 hours and were then used as aged-synchronized colonies (58). These age-synchronized colonies were grown on 5 cm NGM plates seeded with OP50 at 20°C for 96 hours (4 days) prior to behavioral testing.

### Ethanol treatment

Ethanol-treatment plates were prepared as follows (29,33); prior to adding ethanol, NGM plates were seeded with *E. coli* 4 days before the behavioral testing. Ethanol was then infused into the agar by pipetting 4°C cold 100% ethanol onto the agar to the desired concentrations based on the weight of agar in each plate.

Immediately following the addition of ethanol, plates were sealed by parafilm, and incubated at 20°C for at least 2 hours or until the ethanol was absorbed and the surface of the agar was dried. Worms were transferred to ethanol plates 30min prior to the behavioral testing, and the plates were re-sealed with parafilm.

### Behavioral recording

Behavioral recording was done using the Multi-Worm Tracker apparatus and program version 1.2.0.2 (25). Plates containing worms were placed into the Tracker holder on the platform, and visualized using a Falcon 4M30 camera (Dalsa) and a 60mm f-number 4.0 Rodagon (Rodenstock) lens 40 cm above the platform. Images were processed using a capture card PCIe-1427 CameraLink (National Instruments). An elliptical region of interest excluding the outer 5 mm of the tracked plate was denoted on the Tracker program. The recording started immediately after a plate was placed on the Tracker, and worms were given 100s to acclimatize before the first tap stimulus was delivered. Taps were produced by a solenoid tapper that drives a plunger into the side of the plate at a 10s ISI. After 30 taps were delivered to the plate, worm behavior was recorded for 10s more to capture the response to the last stimulus.

### Body curve measurement

To assess body curve data was extracted by the Tracker software “Choreography” as the average angle (in degrees) of the body segments for all qualified worms per frame (mean curve per frame) (25). The mean body curve per worm was calculated from mean curve per frame during the 90s to 95s interval after the recording started (which ended 5s before the first tap stimulation). Student’s t-tests were used to evaluate differences between the average degree of body curve at different ethanol concentrations, with alpha value = 0.05.

### Tap responses analysis

Mean reversal probability, duration and speed of responses occurring within 1s of tap delivery for each plate of worms were calculated using the Choreography program (25). Rapid forward response was defined as a response with forward bias and response velocity larger than the baseline maximum velocity 0.3s to 0.1s before a tap. Velocity was calculated by speed x bias (bias is the movement direction: 1 = forward, -1 = backward, 0 = negligible movement). Matlab® and Python scripts were used to pull and compile raw data from Choreography and then calculate rapid forward response probability from the proportion of worms that respond to a tap within 1s by briefly accelerating forward on a plate. A Pause was defined as a response with forward or reverse baseline bias and a response velocity of 0. A Reversal was defined as a response with pause or forward baseline bias and response velocity smaller than 0. A ‘No Response’ was defined as a response with response velocity within the baseline velocity range and with no change in bias. The probability of a rapid forward response or a reversal response was calculated as the number of worms that had this response divided by the total number of worms that responded to the tap (excluding no response or worms with missing data) for every 0.1 s from 0.1 - 0.5 s after a tap. Plates containing fewer than 10 worms responding to any tap were excluded from this calculation.

### Statistical analysis

The effects of ethanol on habituation in wild-type worms were assessed. “Initial response” was defined as the response for the first tap. “Habituated level” was calculated as the mean response to the last 3 taps (taps 28-30). Student t-tests were used to evaluate differences in initial response and habituated level. Repeated measures ANOVAs with tap as a within subject repeated-measure factor on mean response per plate were used to evaluate the effect of independent variables on tap responses. If more than one independent variable (i.e. strain and dose) was evaluated, then the main effects and the interactions between variables were evaluated. Tukey-Kramer posthoc tests were used to compare habituation curves between groups and to compare responses to individual taps between groups. Each sample (a plate) represents the mean value from a plate containing ∼20-80 worms. “Habituation” was defined as a significant decrease in mean response level of the 28^th^-30^th^ tap compared to the initial response level, evaluated by Tukey-Kramer posthoc tests comparing the difference between the 1^st^ and 28^th^-30^th^ responses. To reduce type I error due to the large sample size (67 independent experiments consist of a total of 229 and 206 plates of worms on 0mM or 400mM ethanol, respectively), alpha was set to 0.001 instead of 0.05.

To assess the impact of genetic manipulations including *slo-1, slo-1::GFP*, or degron strains, repeated measures ANOVAs were used to determine significant effects of ethanol on reversal or rapid forward response probability. For these experiments the critical comparison was the effect of 0 or 400mM ethanol on habituation of response probability within a strain. Tukey’s posthoc test was used to compare these groups. An ethanol group was determined to show significant ethanol effects on reversal or rapid forward response probability if Tukey’s posthoc test showed a significant difference between the 400mM and 0mM control group within a strain.

## Results

### Concentration-dependent effect of ethanol on body curve

To validate our protocol and ensure that our experiments replicated previously reported effects of ethanol on behavior, we assessed worm body curve because ethanol has been reported to flatten worm’s body curve in a concentration-dependent manner (29). The ethanol treatment data collected in this study confirmed this observation (**Fig 1**). Worms exposed to ethanol showed flatter body curves compared to the 0mM control (F(6)=140.68, p<0.001, 0mM vs 100-600mM, all p<0.001; **Fig 1b**). The curvature decreased progressively as ethanol concentration increased from 100mM to 400mM (100mM vs 200-600mM, p<0.001, 200mM vs 300-600mM, p<0.001, 300mM vs 400mM, p=0.012, 300mM vs 500mM, p<0.001, 300mM vs 600mM, p=0.010). However, there were no significant differences in body curve between 400mM and higher ethanol concentrations (400mM vs 500mM or 600mM, p=n.s.). Since increasing the ethanol concentration beyond 400mM did not produce any additional effect on body curve, later experiments in this study used 400mM as the concentration for all ethanol treatments.

**Figure 1:**
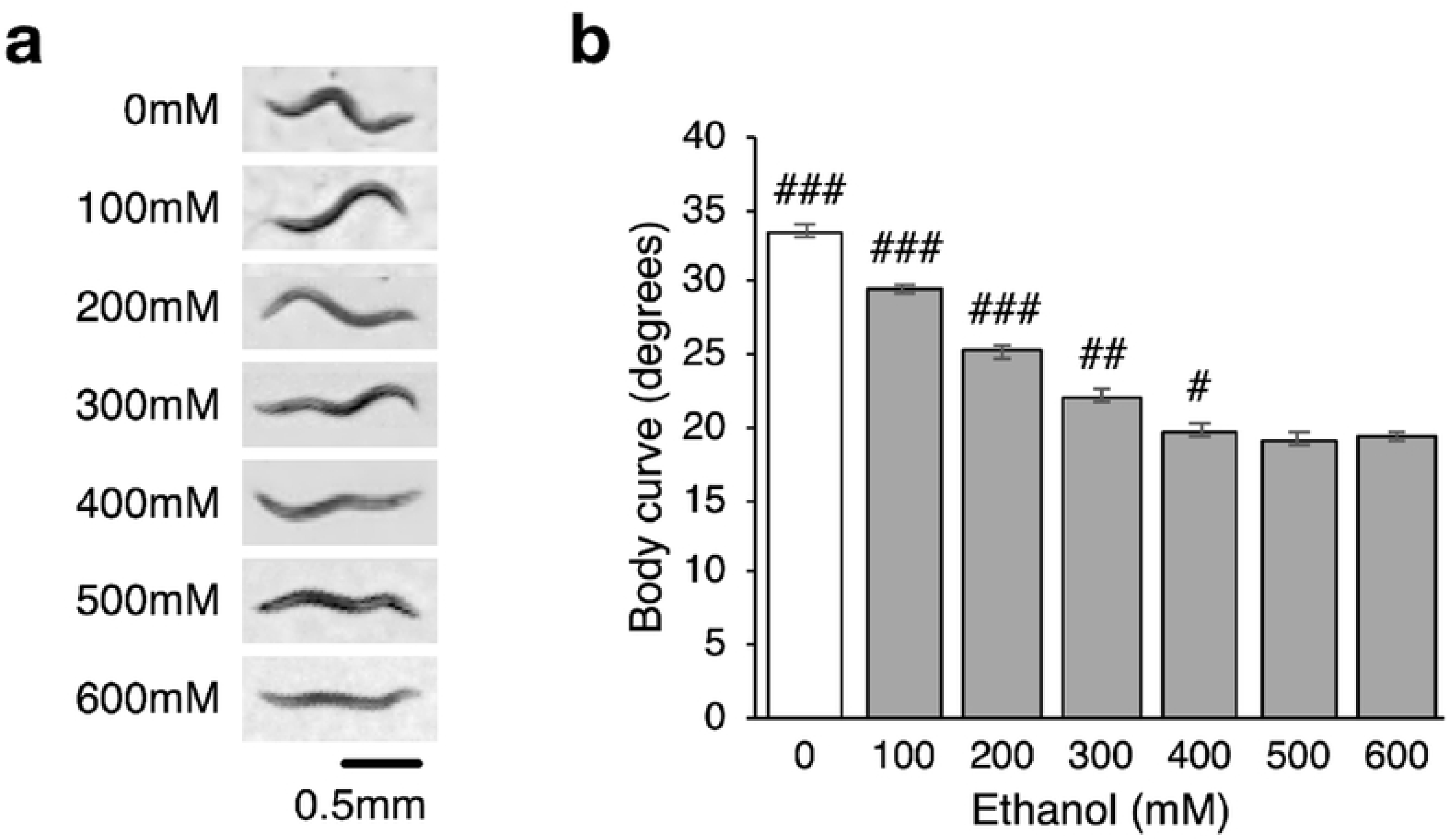
Ethanol flattens the worm’s body curve in a dose-dependent manner. (**a**) Ethanol flattens worm’s body curve progressively as dose increases from 100mM to 600mM. (**b**) Increasing ethanol concentrations significantly reduce the degrees of body curve of worms until the concentration reached 400mM after which further increasing the ethanol concentration had no significant effect on worm body curve. Error bar = SE. # = p < .05, ## = p < .001, ### = p < .0001.

### Ethanol has different effects on habituation of response probability, duration and speed

Having replicated earlier findings on body curve with our protocol and determined that the optimal ethanol dose to use was 400mM, we next investigated the effects of this ethanol dose on habituation to mechanosensory stimuli. Mechanosensory stimuli were delivered by repeatedly tapping the side of an agar filled Petri plate holding the worms mounted on the Multi-Worm Tracker (25). To investigate how habituation of the three main components of the tap reversal response (probability, duration and speed) are affected by ethanol, we conducted a large-scale study of wild-type *C. elegans* that were given 30 repeated mechanosensory stimulations (taps) delivered at 10s ISIs on or off 400mM ethanol.

Our results showed that ethanol produced different effects on the habituation of reversal probability, duration and speed (**Fig 2**). Firstly, ethanol’s effect on initial response level of these three habituation components was different. Initial reversal probability and duration responses of the 400mM ethanol group were significantly reduced compared to the 0mM control group (reversal probability: p<0.001; duration: p<0.001; **Fig 2 a-b**), but initial reversal speed was increased compared to the control group (p<0.001; **Fig 2c**). The final habituated level (the averaged mean responses to the last three taps) for reversal probability was significantly lower in the ethanol group compared to the control group (p<0.001; **Fig 2a**). While the Tukey test comparing final habituated level for reversal duration between the ethanol and control groups had a low p-value (p=0.0027; **Fig 2b**), this was not statistically significant because our alpha value was set at 0.01; therefore ethanol had no significant effect on final level of duration. Interestingly, the effect of ethanol on the final habituated level of reversal speed was the opposite of its effect on reversal probability, as the ethanol group showed a higher final level compared to the control (p<0.001; **Fig 2c**). Finally, the amount of habituation, as measured by the difference between the initial and final response levels, for each of the different response components was also changed by ethanol. Although reversal probability for the ethanol group appeared to habituate more than the no ethanol group (0.47 separated the initial and final levels for the 400mM group vs 0.44 for the 0mM group; p=0.036; **Fig 2a**), this was not statistically significant due to our low alpha. However, for both duration (1.04s for the 400mM group vs 1.53s for the 0mM group; p<0.001; **Fig 2b**) and speed (0.048 mm/s for the 400mM group vs 0.072 mm/s for the 0mM group; p<0.001; **Fig 2c**) the ethanol group did habituate significantly less than the control group. Together, these results confirmed that ethanol affects components of mechanosensory habituation differently in *C. elegans*. Because we observed a large effect of ethanol on both the initial and final level of habituation of response probability, and because a number of environmental factors influence response speed (e.g., temperature, humidity, wetness of the agar plates, depth of *E. coli* lawn) we chose to focus on how ethanol affects response probability for the remainder of this study.

**Figure 2:**
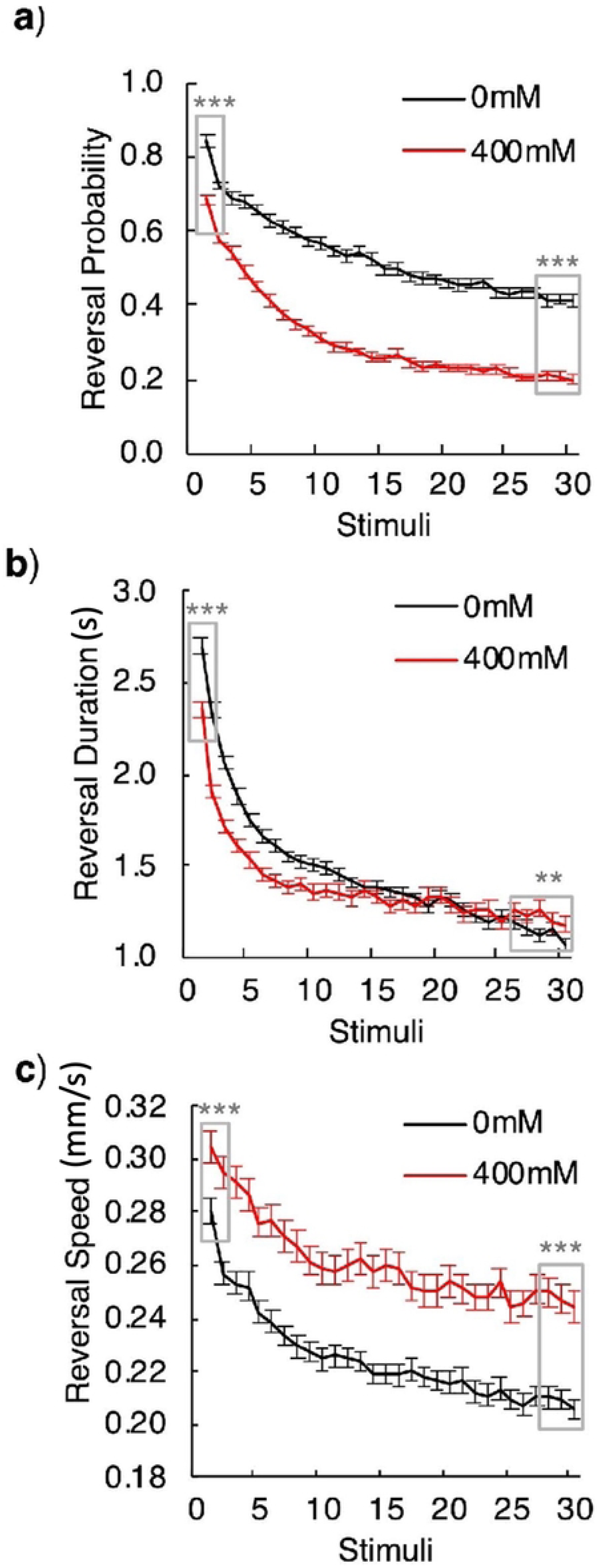
Effects of 400mM ethanol on habituation of tap withdrawal responses in wild-type worms differ by response components. (**a**) Ethanol generally decreased animals’ reversal probability to tap stimuli while not having a large effect on the amount of habituation. (**b**) Ethanol decreased the amount of habituation of reversal duration and (**c**) reversal speed. p < .001. Error bar = SE. * = p < .05, ** = p < .01, *** = p < .001.

### Ethanol changed the dominant response from a reversal to forward movement

Although a reversal is the dominant response to taps for wild-type worms, it is not the only way worms respond to taps (59,60). Worms can also respond to taps by accelerating, pausing, or decelerating. To examine these dynamic responses to taps, changes in speed and direction of movement over time were visualized using direction and speed-based raster plots (**Fig 3a**). For the first tap (Stimulus 1), the raster plots for most of the control group show blue horizontal lines starting from the delivery of the tap stimulus. This corresponds to worms responding to the first tap by reversing and moving backwards for several seconds. Compared to worms on the control plates, the raster plot for the ethanol group shows less blue; this corresponds to the observation that fewer worms on ethanol reversed in response to the first tap (**Fig 3).** From Stimuli 1-5, both the ethanol and control groups showed progressively fewer reversals in response to each stimulus, and fewer worms in the ethanol group reversed in response to taps by stimulus 5 (**Fig 3a**). Raster plots for the last 5 stimuli (Stimuli 26-30) showed shorter and slower reversals and fewer worms responding to the tap in both groups (Fig 3b).

**Figure 3:**
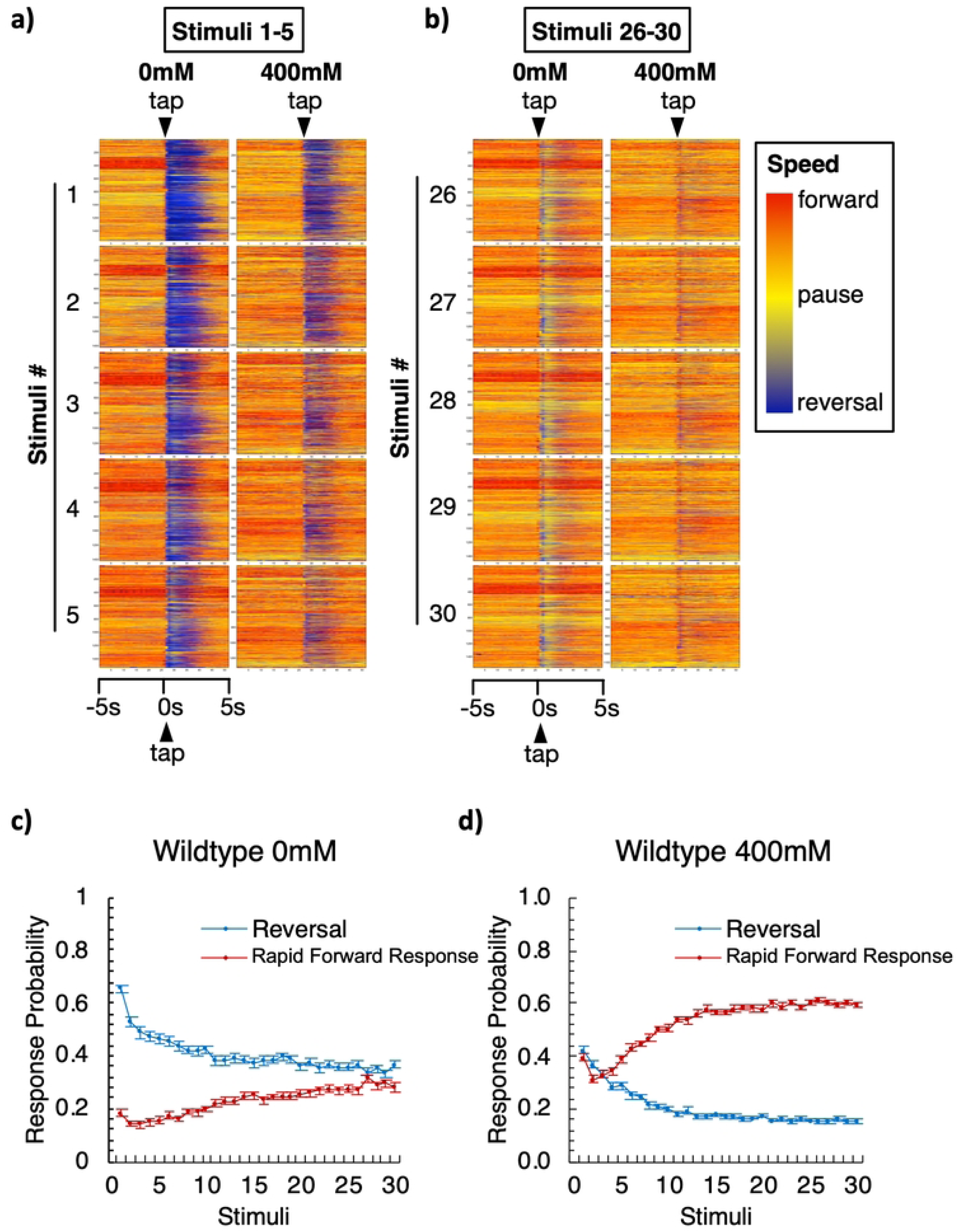
Worms on ethanol are more likely to respond to taps with rapid forward movement than a reversal. (**a-b**) Raster pots of velocity of locomotor movement of individual worms on 0mM or 400mM ethanol in response to the first 5 taps (a) and the last 5 taps (b) delivered at 10s ISI. Within each box, movement of an individual worm (y-axis) across time (x-axis, 5s before and 5s after a tap) is represented as a single horizontal line. The color of the line represents the direction and speed of the worm: the hotter the color, the faster the forward movement (orange to red); the colder the color, the faster the reversal movement (light to dark blue); and pauses are represented by bright yellow. Tap occurs in the middle of each box as indicated by the arrowhead at the top of each column. Data from thousands of worms were stacked vertically to construct a worm speed raster plot (each box). The numbers to the left of each pair of boxes indicates the stimulus number (1-5 or 26-30) in the 10s ISI habituation series on 0mM or 400mM ethanol plates. (**c**-**d**) Percentage of reversal or rapid forward movement response types of the total responses to taps (Stimuli) for wild-type animals (**a**) on 0mM or (**b**) on 400mM ethanol. Errorbar = SE.

When the raster plots for Stimuli 26-30 were examined closely, the response pattern of worms in the ethanol group shows a thin vertical red band at the time of tap delivery (**Fig 3b**). This indicates that to the majority of the worms on ethanol responded to the final stimulus by moving rapidly forward (red) for a very short period of time. In contrast, raster plots for Stimuli 26-30 for the no alcohol group show a thin vertical blue band shortly after the tap was given followed by a wider light yellow vertical band, indicating that instead of long reversals, many worms in the control group responded to the last 5 taps by reversing very briefly then pausing (yellow) for ∼1-2 s. Additionally, some worms simply paused for a few seconds rather than reversing. These data suggest that ethanol does not simply decrease the reversal probability to repeated taps as shown in (**Fig 2a)**, but switched the dominant response direction from a reversal to forward movement.

To quantify this switch in dominant response direction, the probability of reversals or rapid forward responses to each tap were plotted in **Fig 3c-d**. For both groups, the probability of reversal responses decreased as the number of taps increased, similar to data shown in (**Fig 2a)**. In contrast, the probability of rapid forward responses gradually increased as the number of taps increased for both groups (Repeated measures ANOVA: F(58,14268)(response type)=54.083, p<.001; **Fig 3c-d**). For both groups, the ethanol treatment had a significant effect on response type probability (F(24,14628)(dose)=2.33, p<.001, F(58,14268)(response type*dose)=3.16, p<.001). For worms in the no alcohol group, from the first to last tap the probability of a reversal response was always higher than the probability of a rapid forward response (**Fig 3c**). Although the probability of reversal and rapid forward response approached each other during the final taps, at no point was the probability of a rapid forward movement higher than that of reversals (rapid forward vs. reversal responses, tap 22, 27, 29, p=n.s., all other taps, p<.001). In contrast, worms in the ethanol group initially had similar reversal and rapid forward response probabilities (rapid forward vs. reversal responses, tap 1-4, p=n.s.), but after the 4^th^ tap, the rapid forward response probability became significantly higher than reversal response probability (rapid forward vs. reversal responses, tap 5-30, p < .001; **Fig 3d**). These data indicate that ethanol not only altered habituation of reversal responses to taps, but also changed the dominant tap response from a reversal to a rapid forward movement.

### The BK channel ortholog slo-1 mediates ethanol’s effect on tap response plasticity

The BK channel is a well characterized target of ethanol in mammals and in *C. elegans* (29,52,61–65). The BK channel is composed of four pore-forming alpha subunits (**Fig 4a**). Each alpha subunit consists of seven transmembrane domains (S0-S6) and long intracellular hydrophobic segments (S7-10). Within the transmembrane domains, the voltage sensor (S1-4) detects membrane potential changes and the pore-forming domain (S5-6) allows passage of potassium ions. The long intracellular hydrophobic segments (S7-10) contain two main calcium sensors, the RCK1 and RCK2, that allow calcium-coupled channel activation and also contains a putative ethanol binding domain near RCK1 that is thought to be crucial for *slo-1*’s interaction with ethanol (53,65). In *C. elegans*, mutations in this ethanol binding region of *slo-1* block ethanol’s inhibitory effect on crawling speed and egg laying (52). Therefore, we hypothesized that ethanol would have less of an effect on habituation in a *slo-1* mutant with a mutation in this ethanol binding site.

**Figure 4:**
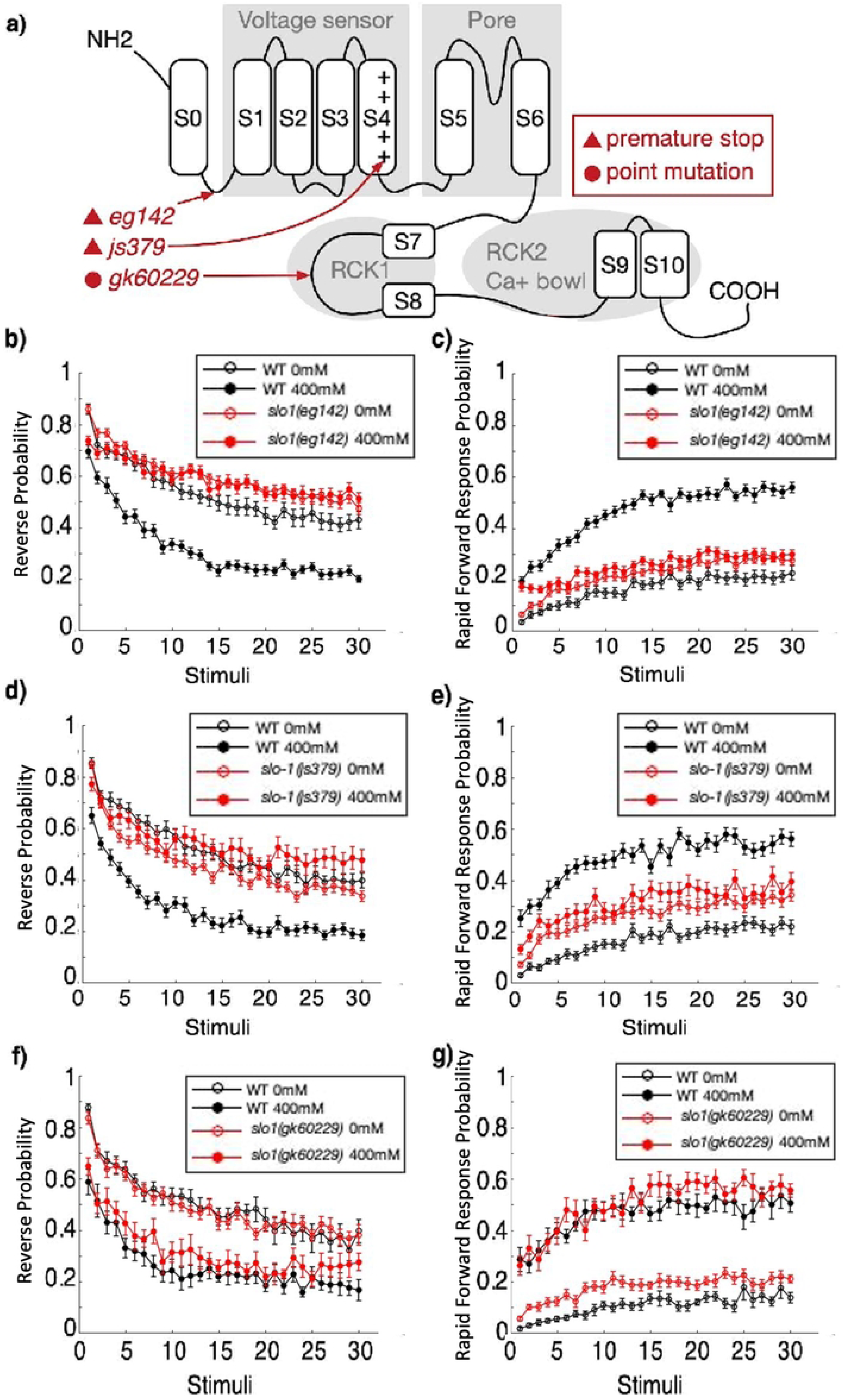
*slo-1* null mutations eliminate ethanol’s effect on reversal and rapid forward response probability while a point mutation in the putative ethanol binding in RCK1 retains the ethanol mediated effect. (**a**) SLO-1 protein structure indicating the locations of three alleles tested by red arrows (66). *slo-1(eg142)* contains a nonsense mutation at the first transmembrane motif S0. *slo-1(js379)* contains a nonsense mutation at the ion pore voltage sensing region. *slo-1(gk602291)* contains a T381I point mutation in the putative ethanol binding region. (**b**) Ethanol did not impact the reversal and (**c**) rapid forward response probability of worms carrying the *slo-1(eg142)* null allele. (**d**) Ethanol’s effect on worms carrying the *slo-1(js379)* null allele is inverted and reduced for reversal probability (**e**) and is lost entirely for rapid forward response probability. (**f**) Ethanol’s effect on worms carrying the *slo-1(gk602291)* allele is the same as on wild-type animals for reversal (**g**) and rapid forward response probability. Errorbar = SE.

To investigate whether the BK channel is involved in the ethanol induced changes in tap response habituation, we tested mutant strains carrying three different *slo-1* alleles (**Fig 4a**). Two of the three alleles, *eg142* and *js379*, are putatively null alleles (29,67) with a nonsense mutation at the first transmembrane motif S0 (66), or at the ion pore voltage sensing region, respectively. The other allele, *gk602291,* has a point mutation in the ethanol binding region near calcium sensing domain RCK1. This point mutation substitutes a highly conserved threonine to isoleucine at position 381 of the SLO-1 channel (T381I) (52). *slo-1(gk602291)* mutants have been shown to have strong resistance to the inhibition of locomotor activity and egg laying by ethanol, similar to that of null mutant *slo-1(js379)*, but unlike in the null mutant basal behaviors such as neck posture (body curve measure of the anterior part of the animal) were unaltered by this mutation (52).

Plates of each of the three *slo-1* mutant strains and wild-type worms were given 30 tap stimuli at a 10s ISI in the presence of either 0mM or 400 mM ethanol. The wild type worms showed large differences for both reversals and rapid forward response when worms in the 0mM groups were compared to the 400 mM groups. In contrast, the 0mM and 400mM groups of worms carrying either of the two null alleles of *slo-1* were very similar for habituation of both reversal or rapid forward movement probability, and both null mutant groups looked similar to the 0mM wild-type worms (**Fig 4b-e**). A repeated measures ANOVA comparing wild-type worms to worms carrying the null allele *slo-1(eg142)*, given 30 taps on and off of ethanol, replicated the original observation that ethanol reduced reversal probability and increased rapid forward response probability in wild-type worms in response to taps (rmANOVA(reversal): F(29,4814)(strain*ethanol*tap)=2.835, p<.001, posthoc: wild-type vs. wild-type ethanol, p<.001; rmANOVA(forward): F(29,4843)(strain*ethanol*tap)=5.934, p<.001, posthoc: wild-type vs. wild-type ethanol, p<0.001), while the presence of ethanol failed to show a significant effect on the reversal or forward movement probability of *slo-1(eg142)* worms (posthoc: *slo-1(eg142)* vs. *slo-1(eg142*) ethanol, p=n.s., for both reversal and rapid forward responses; **Fig 4b-c**). This indicates that the *slo-1(eg142)* null mutation eliminated ethanol’s effect on both reversal and rapid forward response probability to repeated taps. For the second putative null allele *slo-1(js379)*, the repeated measures ANOVA showed a significant effect of ethanol on reversal probability (rmANOVA: F(29,4901)(strain*dose*tap)=2.51, p<.001, posthoc: *slo-1(js379)* vs *slo-1(js379)* ethanol, p=.03; **Fig 4d**), and failed to show a significant effect of ethanol on rapid forward response probability (posthoc: *slo-1(js379)* vs *slo-1(js379)* ethanol, p=n.s.; **Fig 4e**). Interestingly, the effect of ethanol on reversal probability was opposite to that of wild-type, where *slo-1(js379)* mutants showed a slight increase in reversal probability (**Fig 4d**). Importantly, posthoc comparison showed that the reversal probability habituation curve of *slo-1(js379)* mutants for the ethanol or no ethanol groups were not significantly different from that of wild-type, indicating that ethanol had much less of an effect on reversal probability response of the *slo-1(js379)* mutants than on wild-type worms (posthoc: wild-type vs *slo-1(js379)* and wild-type vs *slo-1(js379)* ethanol, p=n.s.). These findings are very similar to the findings from the other null mutation, *slo-1(eg142)*, supporting the hypothesis that *slo-1* plays an important role in how ethanol alters habituation of behavioral responses to repeated stimuli.

Worms carrying the *slo-1(gk602291)* allele with a mutation in the ethanol binding domain have been reported to show similar resistance to ethanol’s inhibitory effect on egg laying and locomotor activity as worms carrying the null allele *slo-1(js379)* (52). However, we observed no significant differences in habituation of reversal and rapid forward response probability between *slo-1(gk602291)* mutants and wild-type worms (rmANOVA(reversal): F(29,1450)(strain*dose*tap)=0.88, p=n.s.; posthoc: *slo-1(gk602291)* vs *slo-1(gk602291)* ethanol, p=<0.001; wild-type vs wild-type, p=<0.001; **Fig 4 f-g**). This suggests that the *slo-1* T381 residue in the ethanol binding domain is not involved in ethanol’s effect on habituation of reversal or rapid forward response probability to repeated taps.

### Cell-specific degradation of the BK channel revealed slo-1 expression in neurons is responsible for the effect of ethanol on response plasticity to repeated taps

The effects of ethanol on habituation might be mediated by ethanol’s effects on the whole animal, on the nervous system, on the muscles or on a specific subset of neurons*. slo-1* is known to be expressed in both neurons and muscles. To investigate which cells are responsible for the role of *slo-1* in mediating the effects of ethanol on habituation, SLO-1 was selectively degraded in all neurons, or specifically in mechanosensory neurons. To do this, worms with GFP inserted into *slo-1*’s C-terminus were crossed into degron strains that drove degradation specifically pan-neuronally (*rgef-1p*) or in the mechanosensory neurons (*mec-18p*).

As a control we first showed that the GFP insertion did not greatly disrupt *slo-1* function during habituation. While there were minor difference between the *slo-1::GFP* strain and wild-type worms (rmANOVA(reversal): F(29,7192)(strain*tap)=2.82, p= <0.001; posthoc: *slo-1::GFP* vs wild-type, p=<0.001), critically ethanol has a significant effect on both reversal and forward movement response probability of *slo-1::GFP* worms (posthoc: *slo-1::GFP* vs *slo-1::GFP* ethanol, p=< 0.001; **Fig 5a-b**). These results suggest that these *slo-1::GFP* worms may have slightly altered baseline habituation responses to repeated taps because of the GFP addition onto SLO-1, or because of differences in the background of the *slo-1::GFP* strain and our wild-type controls. However, these data showed that ethanol’s effect on habituation of reversal and rapid forward response probability in *slo-1::GFP* worms is intact in these worms.

**Figure 5:**
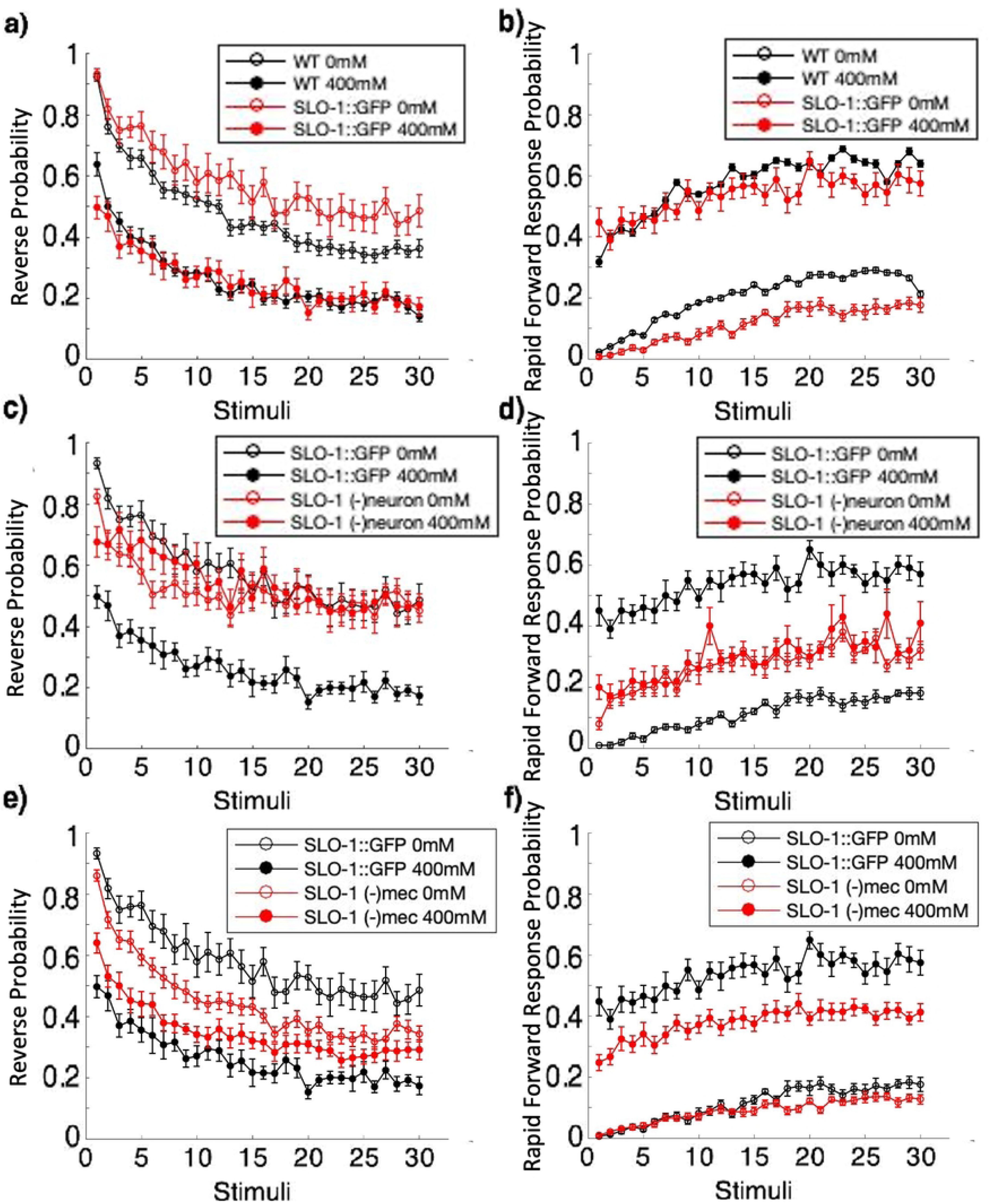
Cell-specific SLO-1 degradation indicates SLO-1 in neurons are associated is required for the effect of ethanol on reversal and rapid forward response probability. (**a**) Ethanol has a similar effect on reversal (**b**) and rapid forward response probability in worms expressing SLO-1::GFP as it does in wild-type worms. (**c**) The effects of ethanol on reversal probability are lost in worms with SLO-1 degraded in neurons (SLO-1 (-)neuron), (**e**) and dampened in worms with SLO-1 degraded in mechanosensory neurons (SLO-1 (-)mec). (**d**) Similarly, the effects of ethanol on rapid forward response probability are lost in worms with SLO-1 degraded in neurons (SLO-1 (-)neuron) (**f**) and in worms with SLO-1 degraded in mechanosensory neurons (SLO-1 (-)mec). Errorbar = SE.

To test whether *slo-1* expression in neurons is responsible for ethanol’s effects on reversal and rapid forward response probability, SLO-1::GFP worms were crossed into a degron strain that targeted degradation of SLO-1 pan-neuronally (SLO-1(-)neuron) and these worms were habituated on and off of ethanol. Results showed that worms without SLO-1 in their neurons failed to show an effect of ethanol on reversal or rapid forward response probability (posthoc(reversal): SLO-1(-)neuron vs SLO-1(-)neuron ethanol, p=n.s.; posthoc(forward): SLO-1(-)neuron vs SLO-1(-)neuron ethanol, p=n.s.; **Fig 5 c-d**). This indicates that *slo-1* expressions in neurons is required for ethanol to affect the plasticity of reversal and rapid forward response probability.

When SLO-1::GFP was degraded in just the mechanosensory neurons, ethanol still had significant effect on reversal probability (posthoc: SLO-1(-)mec vs SLO-1(-)mec ethanol, p=.021; **Fig 5 e-f**). However, the effect of ethanol was smaller than the effects observed in wild-type worms. Similarly, although ethanol did effect rapid forward response probability of SLO-1(-)mec worms (SLO-1(-)mec vs SLO-1(-)mec ethanol, p<.001), and SLO-1(-)mec worms displayed similar rapid forward response probability to the *slo-1::GFP* control when not on ethanol (posthoc: SLO-1(-)mec vs *slo-1::GFP*, p=n.s.), when exposed to ethanol SLO-1(-)mec worms had significantly different rapid forward response probability curves compared to the *slo-1::GFP* control (posthoc: SLO-1(-)mec ethanol vs *slo-1::GFP* ethanol, p<.001). This suggests that while SLO-1 in mechanosensory neurons play some role in mediating ethanol’s effect on plasticity of reversal and rapid forward response probability, SLO-1 in other neurons is likely playing a role as well.

## Discussion

This study used *C. elegans* to examine the effects of ethanol on mechanosensory habituation. We found that ethanol had divergent effects on different response components (probability, duration, and speed), and on different types of responses (reversal vs. rapid forward movement). Exposure to ethanol resulted in a decrease in the initial response for reversal probability and duration but increased it for reversal speed. Ethanol also decreased the final level of response after tap habituation for reversal probability and increased it for reversal speed. Additionally, ethanol decreased the amount of habituation observed for reversal duration and reversal speed and had little effect on the amount of habituation of reversal probability; the ethanol group actually displayed a non-significant increase in habituation of reversal probability. However, while exposure to ethanol did result in a general decrease in reversal probability to the tap stimuli, it simultaneously produced an increase in the probability of a rapid forward response compared to worms in the ethanol free condition. The BK potassium channel SLO-1 was critical for ethanol’s effects on habituation of response probability as null alleles of this gene eliminated ethanol’s impact. Interestingly, a mutation in an ethanol binding domain near RCK1 did not disrupt the effect of ethanol on habituation of response probability, implying that this binding site is likely not involved in ethanol’s effects on this aspect of behavior. Preliminary investigations into where SLO-1 is needed for its effect on habituation suggested ethanol affects habituation of response probability through *slo-1* expression in neurons. As degrading SLO-1 in mechanosensory neurons attenuated, but did not eliminate, ethanol’s effect, SLO-1’s function is probably distributed across the neural circuit that controls mechanosensory responses.

### Ethanol can both facilitate and inhibit habituation

From the first report on alcohol’s effect on habituation in 1967 (13), there has been an ongoing debate about whether alcohol inhibits or facilitates habituation (13–20,68,69). Prior to this report, six publications support an inhibitory effect of alcohol on habituation (13,14,16–18,68), and three publications support a facilitative effect (19,69,70). Only one report encouraged the field to consider a more complex effect of alcohol on habituation than a straight-forward inhibitory or facilitatory role (15). Peeke *et al*. (15) found 0.3% ethanol facilitated habituation of aggressive display frequency towards conspecific male cichlid fish, but inhibited habituation of aggressive display duration (15). However, perhaps due to small sample size (N=7), the authors of this study (15) did not make a strong claim about the bi-directional effects of alcohol on habituation. Although alcohol has since been known to have bi-directional effects on other types of behaviors, there was no strong evidence demonstrating ethanol could have opposite effects on habituation within the same response. Our data, based on a large dataset with high statistical power (77 experiments containing more than 21,000 animals), provides the first convincing data supporting the hypothesis that alcohol can have divergent effects on habituation of different components within the same response (**Error! Reference source not found.**). This work suggests that it is not useful to think of alcohol as a general inhibitor or facilitator of habituation, but rather as a complex modulator of habituation. This would explain why previous findings on ethanol’s impact on habituation were contradictory, as habituation studies typically scored habituation of a single component of a response.

### Ethanol exposure altered response direction from reversals to rapid forward movement

Another significant discovery of this study is that alcohol altered response propensity from reversal to forward movement (**Fig 3**). The significance of this finding can be better appreciated given more background on the natural adaptive behavior in worms. The directionality of worm movement in response to environmental cues can be generally divided into decisions between “avoid” or “approach” (Reviewed in (71)). When given noxious stimuli, such as heat, light, noxious chemicals, high osmolality, or a nose/head touch, worms avoid the stimuli by moving in a direction opposite to the source of the stimuli. Tap stimuli, on the other hand, produce vibrations without a clear direction of the source. In this case, adult worm’s neuronal circuits are biased to produce reversals followed by a change in direction and then forward movement (72–74), suggesting that a reversal followed by a turn is an adaptive response when the direction of the noxious stimuli isn’t clear. In contrast, worms move forward from a tail touch or a heat stimulus to the tail, or to a noxious compound behind the animal; this is most often a forward movement lasting at least several seconds. The brief forward movement we observed in this study does not resemble the natural “approach” behavior. Qualitatively, during approach behavior worms crawl towards the attractive cue, which is accomplished by inhibiting reversals and steering towards the source of the cue (71). This is very different from the brief, short forward movement observed in worms on ethanol in response to tap. These forward responses were as rapid as the reversal responses (both can achieve a speed of 0.3-6 mm/s), and are two times faster than the exploratory/approaching behavior (typically between of 0.1-0.3mm/s) (75). This suggests that the forward response to taps, just like the reversal to taps, might appropriately be described as a “startle” behavior, or could be classified as an “avoid” behavior from the Faumont review (71).

We found that worms on alcohol were equally likely to move rapidly forward or to reverse in response to the first tap, while worms without alcohol were about three times more likely to reverse than to move forward (**Fig 3**). This difference suggests that alcohol disturbed the innate bias to reverse and then change direction when exposed to a stimulus without a clear location. Strikingly, after only a few taps, worms on alcohol became more likely to move forward than to reverse, which never occurred throughout all 30 taps for the control worms not exposed to ethanol. By the 15^th^ tap, worms on alcohol were 3 times more likely to move forward than to reverse. This suggests alcohol produced a behavioral bias opposing the propensity to reverse in response to a startling non-localized stimulus. Previous studies hypothesized that the direction of a worm’s tap response results from the balance of strength between the forward and backward circuit (72,76). Ablation of neurons in the forward circuit increased reversal probability, and ablation of neurons in the backward circuit increased acceleration probability and decreased reversal probability (77). One possibility is that the switch to higher forward movement probability observed in worms on ethanol came from selective inhibition of the backward circuit and/or enhancement of the forward circuit. This altered response bias represents a novel opportunity to further our understanding on alcohol’s effect on behavior.

### The BK channel SLO-1 is necessary for ethanol’s effects on habituation of reversals and rapid forward response probability, however the known ethanol binding site on SLO-1 is not required

This study provided the first evidence that the BK channel is involved in alcohol’s effect on a learning behavior. However, the BK channel’s known ethanol binding domain near RCK1 was not important in mediating this effect on habituation, which is a stark difference from what has been described for other behaviors.

The BK channel’s role in mediating the effects of ethanol on behavior has been well established in *C. elegans* research (29). In fact, *C. elegans* provided the first evidence linking the physiological effect of ethanol on ion channels with its effects on behavior(29). Following that, an ethanol binding site close to the BK channel’s intracellular RCK1 domain was identified (50,53). The BK channel’s role in ethanol modulated behavior in *C. elegans* has since been well characterized, including inhibition of locomotor, egg laying (29), inhibition of pharyngeal activities (64), and a role in acute ethanol tolerance (63). The work reported here demonstrated that SLO-1 also plays a crucial role in ethanol-altered plasticity of responses to repeated stimuli. In the absence of ethanol, we found no evidence of a role for SLO-1 in response plasticity to repeated taps (**Fig 4b-e**). However, null mutations in *slo-1* eliminated the effects of ethanol on habituation of reversal and rapid forward response probability (**Fig 4b-e**), indicating that the BK channel is necessary for ethanol to alter response plasticity.

We also tested animals with the T381I mutation near the RCK1 calcium sensing domain of SLO-1, which substitutes a highly conserved amino acid. Despite being very close to the RCK1 calcium sensing region of SLO-1, it has been hypothesized that the T381 amino acid might be part of a conserved ethanol binding/activation site on SLO-1 as most of the 20 closest amino acid residues are conserved and mutations in T381 cause ethanol insensitivity in worm and mammalian SLO-1 channels (52). Furthermore, the same conserved location on mammalian SLO-1 has been identified as an ethanol binding site and T381 is thought to be structurally important for ethanol binding (50,53,65). It has been shown that ethanol binding at this site facilitates the natural interaction with calcium that gates the SLO-1 channel and is necessary for ethanol induced increases in SLO-1 channel activity (50,53). Previous research found that the T381I mutation played an important role in the effects of ethanol on behavior, making locomotion and egg laying behaviors resistant to inhibition by ethanol, similar to a *slo-1* null mutant (52). Notably, the SLO-1 channel in T381I mutants is still functional, and for ethanol independent behaviors that were impacted by *slo-1* null mutants, such as neck posture, T381I animals exhibited wild-type behavior (52). Interestingly, we found that the T381I mutation did not impact the effect of ethanol on habituation (**Fig 4 f-g**). This suggests that another site on the SLO-1 channel mediates alcohol’s effect on habituation and that the T381I region is not required for all of ethanol’s behavioral modulations. Therefore, different domains of the BK channel may mediate the effects of ethanol on different behaviors. While there might be another site on the BK channel that ethanol directly interacts with, SLO-1 activity can also be regulated by numerous other mechanisms including calcium influx, alpha subunit alternative splicing, modulation by kinases, and regulation by auxiliary beta subunits, (53,65,78). Further investigation is necessary to confirm that other mutations in this ethanol binding site also fail to stop ethanol’s impact on response plasticity, and to elucidate which other SLO-1 regulatory mechanisms or additional ethanol binding sites are required for ethanol effect on mechanosensory habituation response plasticity.

### The effect of ethanol on response plasticity to repeated mechanosensory stimuli is likely mediated by *slo-1* expression in neurons

*slo-1* is widely expressed in neurons and muscles and acts in muscles at the neuromuscular junction (66). By degrading SLO-1 in neurons, we showed that SLO-1 action in neurons is critical for the effect of ethanol on response plasticity to repeated mechanosensory stimuli (**Fig 5c-d**). We further investigated whether ethanol’s effect on response probability was mediated by *slo-1* expressed specifically in the mechanosensory neurons that respond to the tap. The tap withdrawal neuronal circuit is composed of 5 mechanosensory neurons (ALML/R, PLML/R, and AVM), 5 pairs of premotor interneurons (AVAL/R, AVBL/R, AVDL/R, PVCL/R, and RIM), a pair of harsh touch neurons PVDL/R, and a single proprioception neuron DVA (76,77,79). We selectively degraded SLO-1 in mechanosensory neurons and found that SLO-1 in mechanosensory neurons is only partially responsible for ethanol altering habituation of reversal and rapid forward response probability (**Fig 5e-f**). Given these findings, it is probable that the main sites of ethanol’s action are distributed over multiple neurons of the tap withdrawal circuit.

## Conclusion

In this study, we demonstrated a complex effect of ethanol on different response components of habituation to repeated mechanosensory stimuli. The data suggests that ethanol acts as a complex modulator of habituation and explains why earlier work found both faciliatory and inhibitory effects of ethanol on habituation. We also identified a shift in behavioral response bias as a result of ethanol intoxication. These findings promote the use of a more detailed characterization of habituation when investigating the effect of ethanol. Exclusively observing how ethanol effects a single response component might miss important differences in other response components and lead to incomplete conclusions.

We found that the SLO-1 BK channel was required in the nervous system for alteration of habituation of response probability by ethanol, although SLO-1’s known ethanol binding site was found to not be necessary for ethanol’s modulation of habituation of response probability. Future investigation into how ethanol interacts with SLO-1 to mediate changes in response probability could identify a novel ethanol binding site on SLO-1 or an ethanol mediated indirect signaling pathway, advancing our understanding of alcohol’s complex interactions on the nervous system.

## Acknowledgements

We thank Dr. Jon Pierce-Shimomura for generously sharing *C. elegans* strains with us and providing helpful feedback on our manuscript. We also thank Dr. Hongkyun Kim for generously giving *C. elegans* strains to us. Additionally, this work was funded by a Discovery Grant (RGPIN-05558) from the National Science and Engineering Research Council of Canada.

## Supporting Information

The supporting data for this manuscript can be found on Open Science Framework at osf.io/7cujp.

